# Tailocin as an alternative weapon to type VI secretion system in bee gut symbionts

**DOI:** 10.64898/2026.10.28.762571

**Authors:** Yulin Song, Nazhin Nikaeen, Nurila Kambar, Korin R. Jones, Madison Wade, Cecilia Leal, Nancy A. Moran

## Abstract

Antagonistic interactions often shape microbial communities, as co-residents compete for resources. One system of interbacterial antagonism involves tailocins: phage tail-like particles produced by some bacteria to kill close relatives. Their roles in natural communities are poorly studied. Here, we identified putative tailocin loci ubiquitously present in genomes of *Snodgrassella* species, core members of the bee gut microbiota. We purified tailocins from culture supernatants of different *Snodgrassella* strains and found that these inhibited growth of multiple *Snodgrassella* strains but not of strains of other species that cooccur in bee guts. Proteomic analyses corroborated the production of tailocins, and cryogenic electron microscopy imaging revealed their morphology and killing activity via membrane disruption. We generated single- and double-gene knockout mutants of a predator *Snodgrassella* strain that naturally possesses both tailocin and a type VI secretion system (T6SS) that was previously shown to function in intraspecies competition in bee guts. These mutants were used in tests for antagonism against a prey *Snodgrassella* strain in vivo. Co-inoculations within gnotobiotic honeybees including visualization within guts showed that both tailocin and T6SS are important for *Snodgrassella* in intraspecies competition. Each of these two antagonism mechanisms inhibits the establishment of susceptible strains within the ileum. Outcomes depend on the timing of colonization by competing strains, with both tailocin and T6SS giving a stronger advantage when the producing strain colonizes first. Thus, tailocin production can underlie priority effects that have been widely observed in the bee gut community and other host-associated bacterial communities.

**Significance Statement:** Interbacterial competition plays a pivotal role in determining bacterial colonization success in natural habitats, including guts of animals. Tailocins are phage tail-like bacteriocins that are widely encoded in bacterial genomes and that function in intraspecies killing. They have been little studied in natural communities, including those in gut environments. Here, we show that bee gut bacteria produce tailocins that kill close relatives within the gut. Tailocins function as phage-derived antagonism mechanisms together with T6SS, which are also phage-derived. The possession of both weapons provides a double safeguard, supporting that intense competition happens in bee gut environments. Our findings bring new insights into roles of antagonistic weapons in natural communities.

## Introduction

In many habitats, genetically diverse bacteria live in dense communities, as in the gut microbiota of animals including humans and honeybees, where bacterial cells in close proximity compete for niches and resources (1-3). Whereas high strain-level diversity is observed across hosts (4-6), one or few strains often dominate within individual hosts (5-11). This strain–strain incompatibility reflects nutrient or interference competition between closely related bacteria (11-14). The outcomes of competition between close relatives are often subject to priority effects in which the earlier arriver has a niche-occupying advantage over the later arriver (15-18). Priority effects can result in stable communities that are resistant to invasion by outside strains (19-21). Antagonistic interactions that inhibit or kill nearby competitors play crucial roles as mechanisms underlying priority effects (22, 23). These interactions may involve the release of diffusible antimicrobials, for example, colibactin and some secreted bacteriocins (24, 25), or direct delivery of toxins via contact-dependent weapons (23, 26). Type VI Secretion Systems (T6SSs) and tailocins are two sophisticated systems that function in interbacterial competition and differ in whether contact is required for killing (27, 28).

The T6SS resembles an intracellular phage tail structure, which consists of a rigid tube tipped by a sharp spike and wrapped with a contractile sheath (29-31). This harpoon-like structure is anchored to the cell envelope by a baseplate and membrane complex (32). The contraction of the sheath enables penetration into the neighboring target cell and delivery of effectors in a contact-dependent manner (31). T6SSs recruit a wide arsenal of cargo effectors that act on a broad range of cellular components (28, 33, 34). Although they can target eukaryotic cells, most identified effectors have antibacterial activity, suggesting a primary ecological role in interbacterial competition (28, 35). They drive competition within host-associated communities (12, 36-40).

Like T6SSs, tailocins also share an evolutionary origin with bacteriophages (41-44). They are induced by mutagens such as UV and mitomycin C (MMC), assembled intracellularly, and released as diffusible killing particles (27, 44). Delivery requires lysis and death of the producing cell, imposing a cost of tailocin production. In contrast to phages, tailocins lack the DNA-containing capsid and genes for head formation, nucleic acid replication and packaging (27, 41). Tail fibers show specificity to cell surface receptors of target bacteria (45-49). Work in *Pseudomonas aeruginosa* identified two canonical types of tailocins, the F and R types (41, 50). The R type contains a covering sheath, the contraction of which causes insertion of the inner tube into the membrane of the target bacterium, leading to depolarization of membrane potential and death of target cell (44, 48, 50-53). For the F type, which has no sheath and is thus non-contractile, the killing mechanism has not been well studied (44, 50).

The Western honeybee, *Apis mellifera*, possesses a simple gut microbiota that contains about eight core bacterial species (2). Among them, *Snodgrassella* has evolved with bee hosts for about 80 million years (54). It colonizes the bee ileum, with high diversity in strains and gene repertoires (55). Several other species-level clades of *Snodgrassella* occur in other bee species (56). Two types of T6SS loci are frequently present in *Snodgrassella* genomes (38). Previous work demonstrated that one type (T6SS-1) plays a role in intraspecies antagonism and the other type (T6SS-2) affects host immune responses (39). This study focuses exclusively on T6SS-1, hereafter referred to as T6SS. Other antagonism mechanisms have not been studied in *Snodgrassella* or other community members.

In this study, we identified putative tailocin loci in *Snodgrassella* strains, purified tailocins from cultures, examined them via proteomics and cryogenic electron microscopy (cryo-EM) imaging, and carried out in vitro inhibition assays demonstrating their activity and target range. Using gnotobiotic honeybees with defined inoculations (57), we then investigated the role of tailocin in interbacterial competition in bee guts and its interplay with T6SS.

## Results

### Putative tailocin loci are widespread in *Snodgrassella* genomes

We collected all available genome sequences of *Snodgrassella* isolates, including *S. alvi* from *A. mellifera* and other *Snodgrassella* spp. from other honeybees (*Apis* spp.) or bumblebees (*Bombus* spp.). We identified phage-like sequences and clustered them based on sequence distance (see details in SI Appendix, SI text). Phage-like loci were found in all 83 *Snodgrassella* genomes and form 5 major clusters based on pairwise k-mer Mash distances and the elbow method (58, 59) (Fig. 1A and Fig. S1A). Four of these clusters appear to be prophages, as they possess loci encoding relatively complete phage structural and enzymatic proteins.

**Figure 1.**
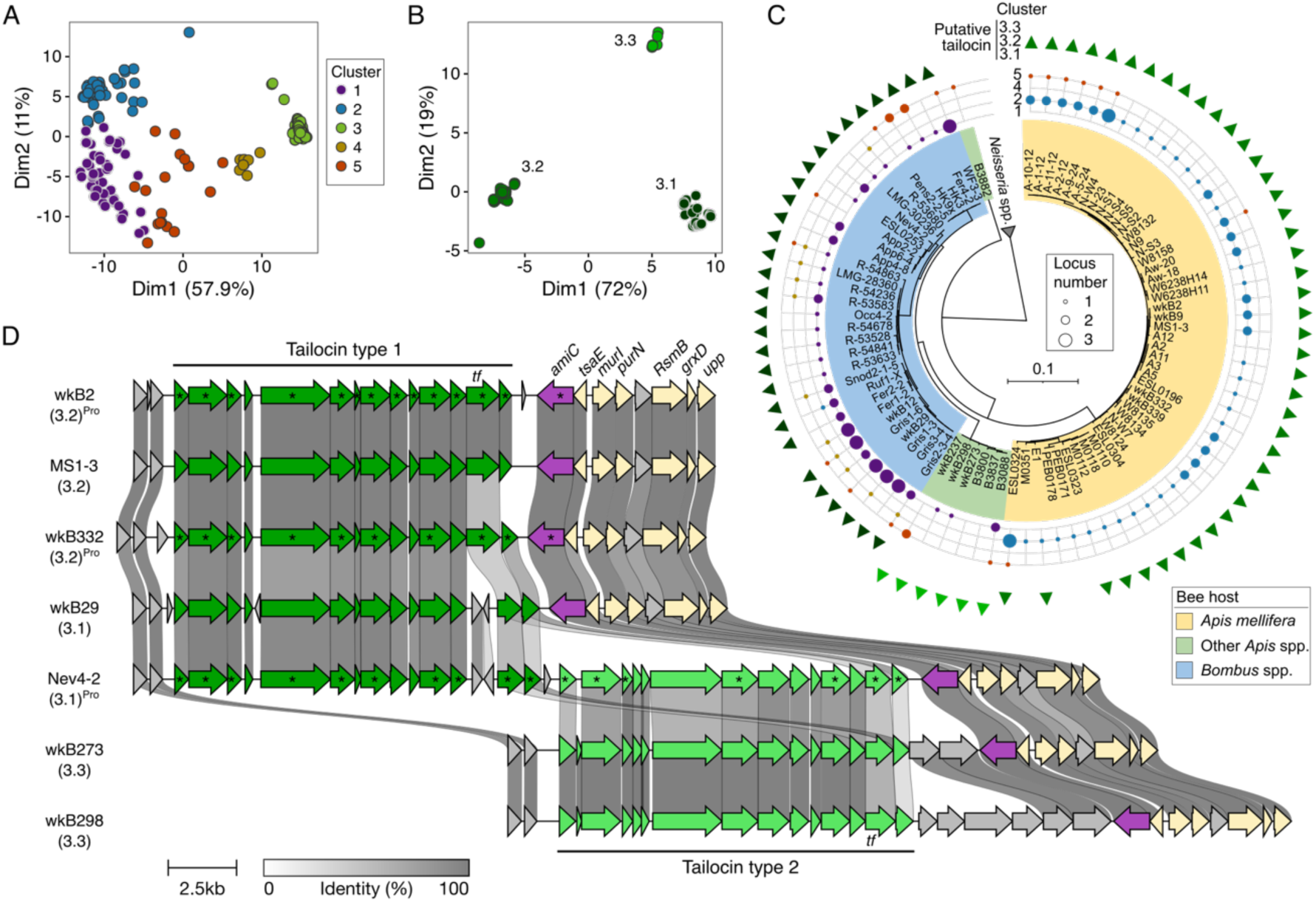
Clustering and annotation of phage-like loci identified in *Snodgrassella* genomes. (A) *Snodgrassella* phage-like loci can be clustered into five major clusters based on pairwise k-mer Mash distances and the elbow method. (B) The putative tailocin cluster (c3) can be further clustered into three subclusters. (C) Distribution of different (sub)clusters of phage-like loci in *Snodgrassella* genomes. The maximum-likelihood phylogenetic tree is based on single-copy genes of the genomes. The outer triangles show the presence of predicted tailocin locus. Colors of (sub)clusters are corresponding to A and B. (D) Synteny of putative tailocin loci from genomes of representative strains. Subclusters are in parathesis. Strains labeled with “^Pro^” were analyzed using proteomics; asterisk-marked genes were detected to be expressed. The two different types of putative tailocins are in dark and light green. The lysin gene is in purple. Homologous genes are linked, with greyscale reflecting the level of amino acid identity. Gene abbreviations and products: *tf*, tail fiber; *amiC*, *N*-acetylmuramoyl-L-alanine amidase (MurNAc-LAA); *tsaE*, tRNA (adenosine(37)-*N*6)-threonylcarbamoyltransferase complex ATPase subunit type 1; *murI*, glutamate racemase; *purN*, phosphoribosylglycinamide formyltransferase; *RsmB*, NOP family class I SAM-dependent RNA methyltransferase; *grxD*, Grx4 family monothiol glutaredoxin; *upp*, uracil phosphoribosyltransferase.

The remaining cluster (c3) encodes tail, baseplate, and lysin proteins but no head proteins or enzymes for replicating and packing. We hypothesized that this cluster encodes tailocins. This cluster forms three subclusters (Fig. 1B and Fig. S1B). The distribution of the prophage-like clusters is correlated to genomic phylogeny and host groups (Fig. 1C). For example, c1 is present in *Bombus* spp. and other *Apis* spp. but not in *A. mellifera*, while c2 is present only in *Apis* spp. The tailocin cluster, however, is present in almost all genomes (78/83, 94.0%), and the distribution of the subclusters reflects phylogeny of the bacterial hosts, as inferred on the basis of single copy genes shared across *Snodgrassella* genomes. The three subclusters exhibit conserved organization and are always flanked by the same set of neighboring genes (Fig. 1D). Subclusters c3.1 and c3.2 share the same type of tailocin gene cluster, whereas c3.3 has a different type. Some bumblebee-associated strains have two types. For example, strain Nev4-2 has a tailocin locus showing high sequence homology to other c3.1 loci but also an insertion that has relatively low homology with those of c3.3, suggesting a lateral transfer event. The same lysin gene, encoding a *N*-acetylmuramoyl-L-alanine amidase (MurNAc-LAA), is shared and highly conserved across all tailocin loci. The two types of gene clusters encode independent tail fiber genes. Notably, the tail fiber genes of different gene clusters show lower sequence identity as compared to other components.

### Purified samples showed tailocin-dependent in vitro inhibition activity

Tailocins can be released into spent medium after mutagen induction (27, 44). To characterize putative tailocins of *S. alvi*, we purified proteinaceous samples from the supernatant of UV-treated cultures of *S. alvi* wkB2^T^ which encodes a c3.2 tailocin (Fig. 2A). SDS-PAGE analysis revealed four major bands of about 38, 48, 95, and 300 kDa (Fig. 2B). By spotting the purified sample onto agar plates with background growth of different bee gut symbiont strains, we found that the wkB2-derived sample inhibits multiple other *S. alvi* strains (Fig. 2C). In contrast, a tail fiber deletion mutant, Δ*tf*, lost the killing capacity. Then, we purified samples from multiple “predator” *Snodgrassella* strains possessing putative tailocin loci and tested inhibition against a list of “prey” strains from different bee hosts. Samples purified from strains wkB2 and MS1-3 (99.1% average nucleotide identity) showed the same inhibition spectrum that is different from those of the more distant strains, wkB273 and Nev4-2 (Fig. 2D). We also tested inhibition against other core species that co-occur in the honeybee gut, including *Gilliamella apicola, Gilliamella apis, Frischella perrara*, *Bartonella apis*, *Bifidobacterium asteroids*, *Lactobacillus helsingborgensis*, and *Bombilactobacillus mellifer* (Table S1). None of these other species was inhibited by wkB2 tailocin. To understand whether mutagen induction is required for *S. alvi* tailocin production, we purified supernatant samples from wkB2 culture induced by UV, MMC, or not induced and tested inhibition against wkB332 target (Fig. S2). To our surprise, non-induced sample showed similar levels of inhibition against the target strain as the exposed samples, indicating that tailocin is constitutively produced without induction.

**Figure 2.**
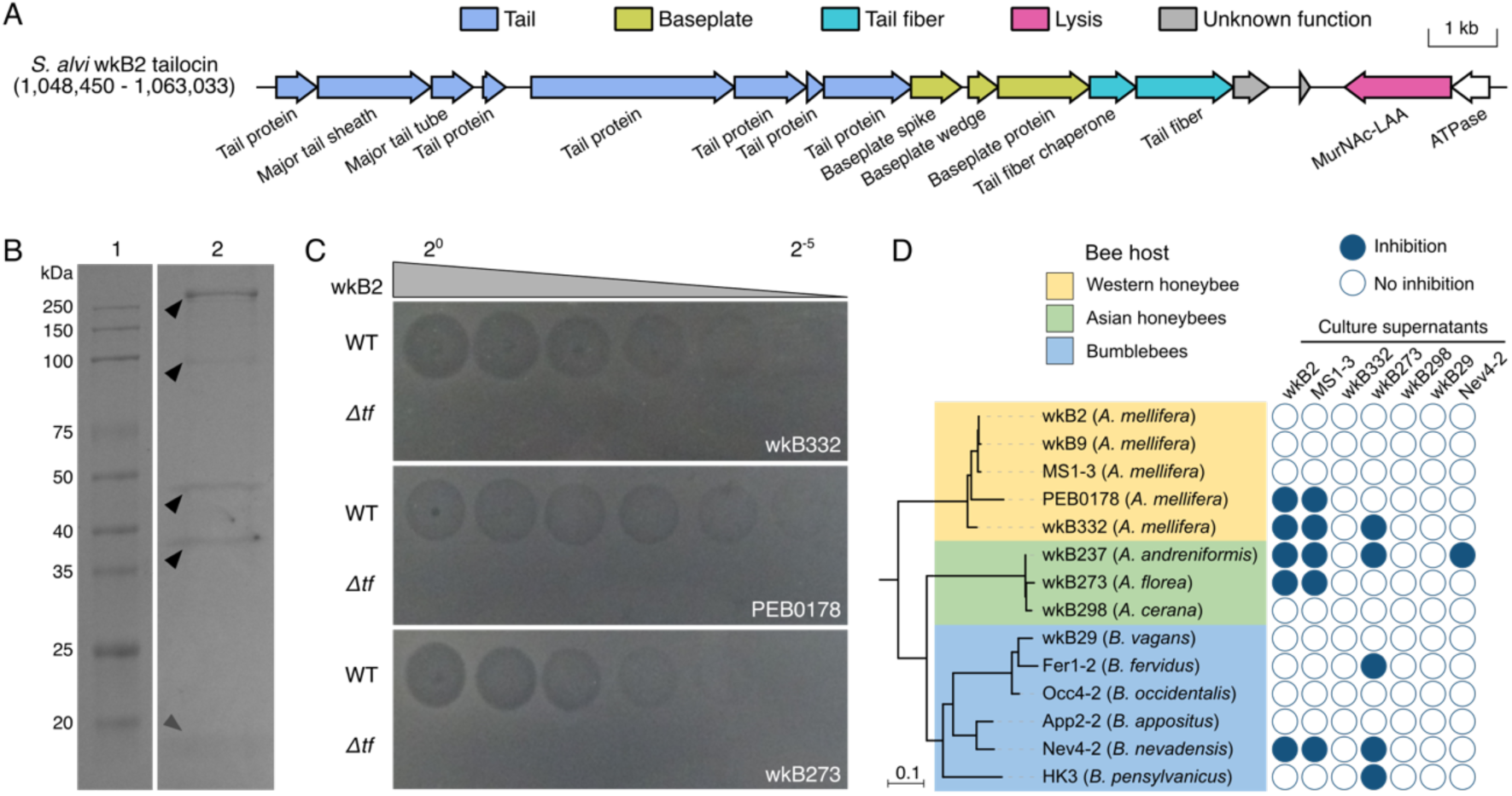
*Snodgrassella* strains produce tailocins inhibiting close relatives. (A) Genomic annotation of the tailocin locus in *S. alvi* wkB2 (GenBank: CP091515). (B) SDS-PAGE analysis of the tailocin-containing sample purified from wkB2 culture supernatant. Black triangles indicate characteristic bands; grey arrow indicates another band that is either characteristic or from degradation. (C) Inhibition assay of samples purified from wkB2 WT or Δ*tf*. Prey strains were grown on the background, and serially diluted samples were spotted on the top. (D) Inhibition of samples purified from multiple potential predator strains against an array of prey strains from different bee hosts.

### Proteomics and cryo-EM revealed the production and killing activity of tailocins

To examine the production of tailocin proteins, we performed proteomic analysis on supernatant samples purified from different strains (Fig. 1D and Fig. S3). In our first attempt with wkB2 WT, we detected 14 out of the 15 encoding proteins of the tailocin locus. In the second attempt, we tried multiple representative strains, including wkB2 WT and Δ*tf* (9/15 proteins detected for both), wkB332 (10/15), and Nev4-2 (17/28). We detected most tailocin-locus proteins in all assayed strains, which suggest that the loci are actively expressed. The remaining proteins were likely not observed due to the limit of detection, as many detected proteins only have few spectral counts. We also detected tailocin production in the gut of gnotobiotic bees inoculated with wkB2 WT or Δ*tf*, although only 2 tailocin proteins were detected, which is likely due to the limited scale of purification as far fewer wkB2 proteins were detected in gut samples than in pure culture samples.

We performed cryo-EM to gain insights into the morphology and killing mechanism of *S. alvi* tailocins. Tailocins produced by wkB2 are typical R-type cylinder-shaped particles, with a length of 177 nm and a width of 19 nm (Fig. 3A and B). We did not see similar particles in a sample purified from Δ*tf*, suggesting that tailocins are not properly assembled in this mutant. To understand how tailocin functions, we incubated wkB2 tailocins with cells of a prey strain wkB332. We observed disruption of the prey cell membrane by a contracted tailocin (Fig. 3C). Another prey cell exhibited many contracted and uncontracted tailocin particles attached to or in close proximity to its outer membrane (Fig. 3D). Within the same tailocin-exposed sample, we observed a wkB332 cell that retained an intact double-membrane structure with no apparent signs of tailocin-induced damage (Fig. 3E), and a cell that exhibited signs of lysis with disrupted double-membrane cellular morphology, likely caused by tailocin attack (Fig. 3F). In previous studies using the same imaging protocol, but in which *S. alvi* was not exposed to attacking tailocin, these morphological disruptions were not observed (60).

**Figure 3.**
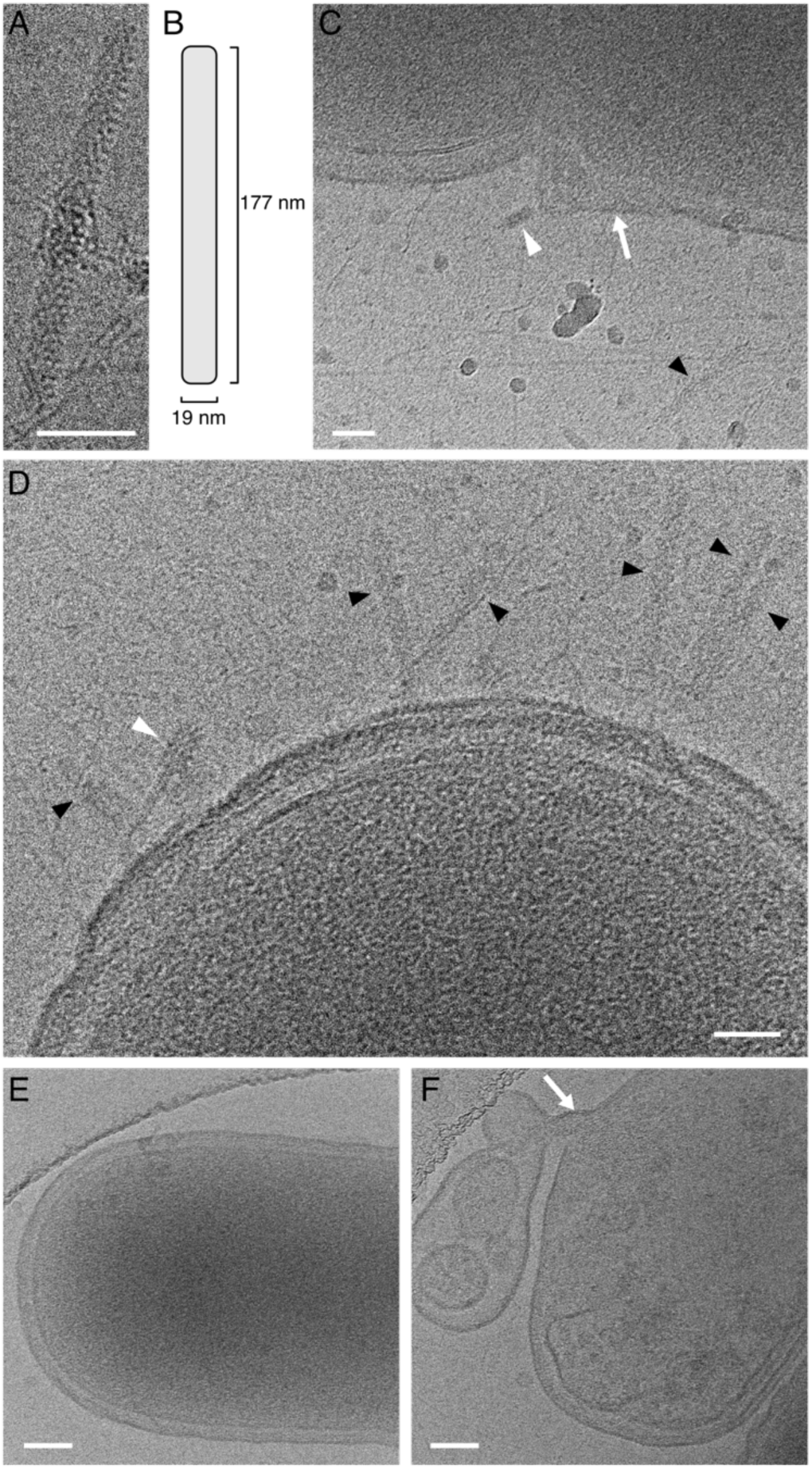
Cryo-EM imaging of *S. alvi* wkB2 tailocins and their interaction with the prey wkB332 strain following incubation. (A) Representative single tailocin particle of wkB2. (B) Schematic illustration of the dimensions of the tailocin particle shown in (A). (C) A triggered tailocin disrupting membrane integrity of a target cell. The white arrow indicates potential tailocin-attacking position and white arrowhead contracted tailocin. (D) Tailocin particles attached to, or located in close proximity to, the surface of a target cell. White and black arrowheads indicate contracted and uncontracted tailocins, respectively. (E) A healthy cell of wkB332. (F) A lysing cell of wkB332 after incubation with wkB2 tailocin. The white arrow indicates a site of membrane disruption characterized by outward bulging and loss of membrane integrity. Scale bars: (A), (C), (D) 50 nm; (E), (F) 100 nm.

### Tailocin and T6SS are both involved in intraspecies competition in vivo

To investigate roles of tailocin and T6SS and their potential interplay in the bee gut, we used strain wkB2 that possesses tailocin and T6SS and strain wkB332 that is susceptible to both tailocin and T6SS (Fig. 4A). For wkB2, besides the wild-type (WT), we included tailocin-null (Δ*tf*), T6SS-null (Δ*tssE1*), and both-null (Δ*tf*Δ*tssE1*) mutants (Fig. 4B). An evolved strain resistant to rifampicin (B332R) was used to distinguish wkB332 from wkB2 strains. Gnotobiotic honeybees were inoculated with one strain or with two strains via oral feeding. In the double inoculations, the two strains were inoculated either simultaneously or sequentially with a 1 d interval (Fig. 4C). Whole guts were dissected after 5 d and plated on medium with antibiotics enabling selection of different strains, and colony-forming units (CFUs) were counted to estimate strain abundance.

**Figure 4.**
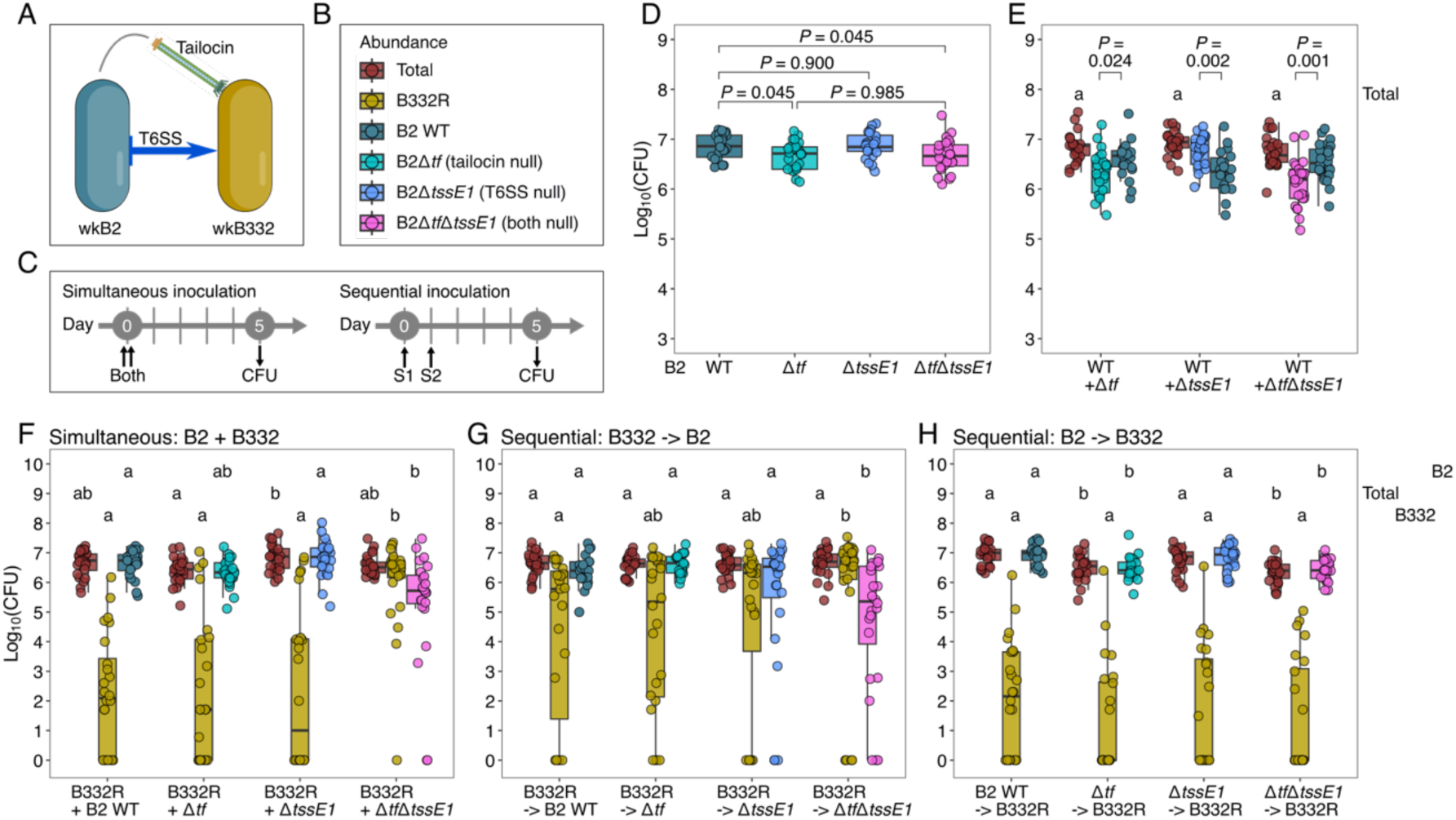
Tailocin and T6SS are involved in intraspecies competition of *Snodgrassella* strains in bee guts. (A) Schematic of the experimental strains. wkB2 possesses tailocin and T6SS; wkB332 is susceptible to both. (B) A key to the strains used in the CFU quantification of strain abundance in D–H. (C) Timelines of simultaneous and sequential oral inoculation of two strains (S1 and S2) in gnotobiotic bees. (D) Mono-colonization of wkB2 WT and tailocin or T6SS gene deletion mutants. Statistical significance was determined by a one-way ANOVA test (*F*(3, 92) = 3.81; *P* = 0.013; *n* = 96 bees) followed by post-hoc pairwise comparisons of estimated marginal means with a Benjamini-Hochberg adjustment. (E) Pairwise simultaneous inoculation of wkB2 WT and each mutant strain. Total bacterial abundance (share the same letter) did not differ significantly among treatment groups, as determined by a one-way ANOVA test (*F*(2, 68) = 2.23; *P* = 0.12; *n* = 95 bees). Within group *P* values were determined by two-tailed *t*-tests (F) Pairwise simultaneous inoculation of wkB332 and each wkB2 strain. (G) Pairwise sequential inoculation of wkB332 followed by each wkB2 strain. (H) Pairwise sequential inoculation of each wkB2 strain followed by wkB332. In F–H, statistical significance was determined by Kruskal–Wallis tests followed by Dunn’s tests with Benjamini–Hochberg method for *P* value adjustment; groups do not share same letters in the same row are significantly different. Each bee represents a biological replicate.

We first compared the colonization levels of the wkB2 strains in mono-inoculations. Δ*tf* and Δ*tf*Δ*tssE1* colonized at lower levels than did WT, while Δ*tssE1* did not differ significantly from the WT (Fig. 4D). When each mutant was simultaneously co-inoculated with the WT, Δ*tf* and Δ*tf*Δ*tssE1* still colonized at lower levels than WT, whereas Δ*tssE1* had an advantage over the WT (Fig. 4E). These results indicate a colonization disadvantage of an intact T6SS, as found previously (61), and an advantage of possessing intact tail fiber.

Next, we co-inoculated each of the wkB2 strains with B332R, simultaneously or sequentially, to test effects of the weapons and inoculation timing (Fig. 4F–H). When simultaneously inoculated, B332R was outcompeted by wkB2 WT, Δ*tf*, and Δ*tssE1*, but not by Δ*tf*Δ*tssE1* (Kruskal–Wallis test on B332R abundance, χ^2^ = 34.0, d.f. = 3, *P* = 1.97e-7, n = 96 bees; *P* < 1e-5 between groups of Δ*tf*Δ*tssE1* and other wkB2 strains, and *P* = 1.0 in other pairwise comparisons). When B332R was inoculated first, its abundance still differs significantly among groups subsequentially inoculated with different wkB2 strains (Kruskal–Wallis on B332R, χ^2^ = 8.99, d.f. = 3, *P* = 0.029, n = 95 bees) and between groups inoculated with wkB2 WT and Δ*tf*Δ*tssE1* (*P* = 0.032). Groups Δ*tf* and Δ*tssE1* show lower median B332R abundance compared to Δ*tf*Δ*tssE1* although they are not statistically different from Δ*tf*Δ*tssE1* or WT. When wkB2 strains were inoculated first, however, B332R abundance was not significantly different among the groups (Kruskal–Wallis on B332R, χ^2^ = 2.98, d.f. = 3, *P* = 0.40, n = 96 bees). Interestingly, the first inoculum of sequential inoculation seems to determine the final total abundance: when B332 was inoculated first, the four groups are not significantly different; however, when wkB2 strains were inoculated first, the tailocin-null groups are significantly lower, following the same pattern of their mono-colonization.

We assessed the same competition data (Fig. 4F–H) using different statistics (Table 1). For each bee, we compared the abundances of wkB2 and B332, to identify how the two antagonism systems and the order of inoculation affect the relative abundances of the two strains. When simultaneously inoculated, wkB2 strains outcompete wkB332, except for wkB2 Δ*tf*Δ*tssE1* which loses its advantage (*P* < 0.001 compared to other groups). When B332R was inoculated first, the competition results show a similar but less pronounced trend: wkB2 strains outcompete B332 except for wkB2 Δ*tf*Δ*tssE1* (*P* < 0.05 compared to other groups). When inoculated first, wkB2 strains, whether WT or mutant, outcompeted B332 in every individual bee.

**Table 1.** Strain dominance in pairwise competition between wkB332 and each wkB2 strain under different inoculation conditions. Raw data are the same as Fig. 4 F–G. A strain wins in the gut of an individual bee when its abundance is higher than that of the other strain. Statistical significance among groups under each condition was determined by Pearson’s Chi-squared tests. Pos-hoc pairwise comparisons between groups were using Fisher’s exact tests. Groups do not share same letters in the same condition are significantly different.

| Inoculation condition | wkB2<br>win | wkB332<br>win | Total | Statistical<br>significance |
| --- | --- | --- | --- | --- |
| Simultaneous: B2 + B332 ($\chi^2 = 42.7$, d.f. = 6, $P = 1.34\text{e-}7$, n = 96 bees) | | | | |
| B332R + B2 WT | 24 | 0 | 24 | a |
| B332R + B2 $\Delta tf$ | 21 | 3 | 24 | a |
| B332R + B2 $\Delta tssE1$ | 22 | 2 | 24 | a |
| B332R + B2 $\Delta tf \Delta tssE1$ | 7 | 17 | 24 | b |
| Sequential: B332 -> B2 ($\chi^2 = 13.3$, d.f. = 6, $P = 0.038$, n = 95 bees) | | | | |
| B332R -> B2 WT | 12 | 11 | 23 | a |
| B332R -> B2 $\Delta tf$ | 16 | 8 | 24 | a |
| B332R -> B2 $\Delta tssE1$ | 13 | 11 | 24 | a |
| B332R -> B2 $\Delta tf \Delta tssE1$ | 4 | 20 | 24 | b |
| Sequential: B2 -> B332 (NA, n = 96 bees)<sup>a</sup> |  |  |  |  |
| B2 WT -> B332R | 24 | 0 | 24 | - |
| B2 $\Delta tf$ -> B332R | 24 | 0 | 24 | - |
| B2 $\Delta tssE1$ -> B332R | 24 | 0 | 24 | - |
| B2 $\Delta tf \Delta tssE1$ -> B332R | 24 | 0 | 24 | - |

<sup>a</sup> Statistical analysis is not applicable (NA) as all groups have identical values.

To understand the spatial context of competing strains in the bee ileum, we performed fluorescence microscopy to directly visualize their colonization. wkB332 and wkB2 strains were transformed with plasmids expressing green fluorescent protein (GFP) and E2-crimson (E2C), respectively. When mono-inoculated, wkB332 colonized the whole ileum, with a strong preference to aggregate at the anterior end (Fig. 5A). The simultaneous inoculation of wkB2 WT, Δ*tf*, or Δ*tssE1*, largely inhibited the establishment of wkB332 in the ileum (Fig. 5B–D). For wkB2 Δ*tf*Δ*tssE1*, however, wkB332 colonized well along the length of the ileum, with a more even distribution compared to mono-colonization (Fig. 5E).

**Figure 5.**
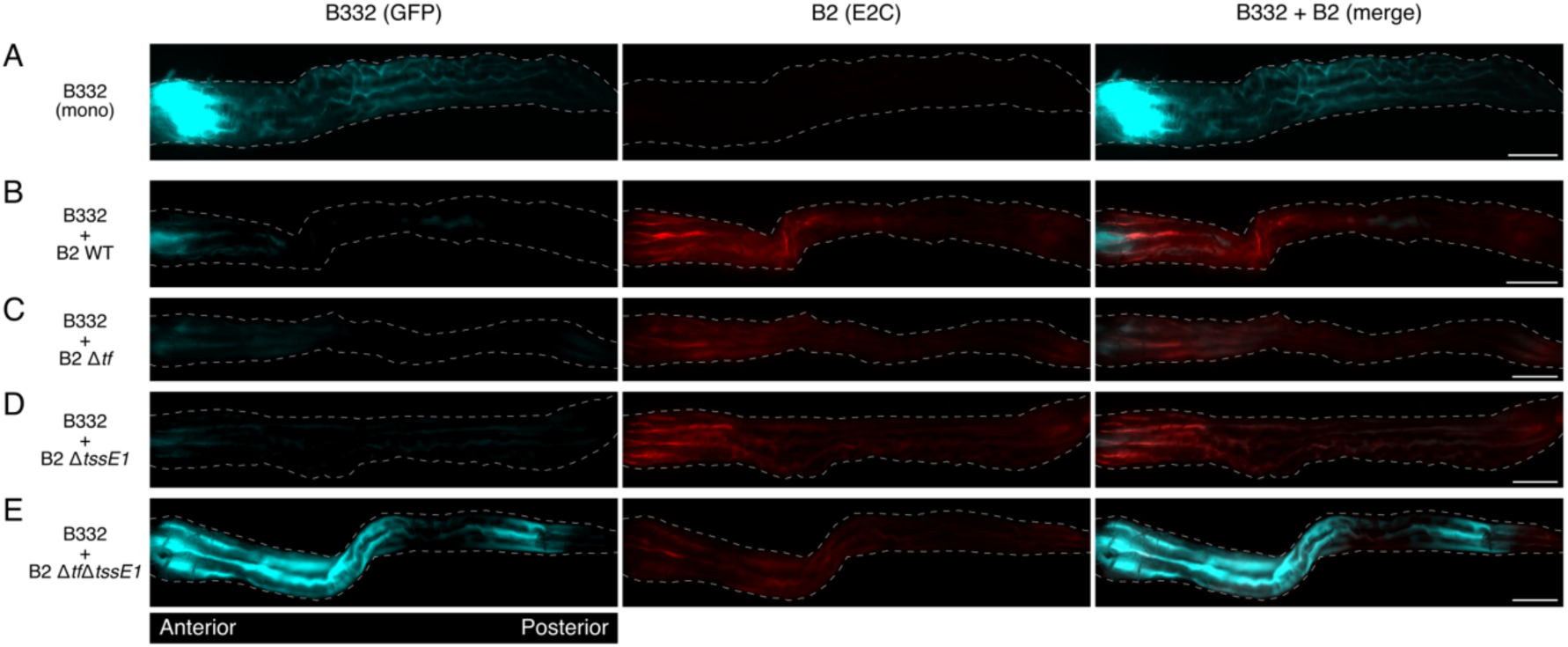
Fluorescence microscopy showing the colonization of wkB332 and wkB2 WT/mutants in the bee ileum. Strain wkB332 expressing GFP and each wkB2 strain expressing E2C were simultaneously inoculated into gnotobiotic bees, and fluorescence was checked after 5 d. (A) Mono-colonization of wkB332. (B) Simultaneous colonization of wkB332 and wkB2 WT. (C) Simultaneous colonization of wkB332 and wkB2 Δ*tf*. (D) Simultaneous colonization of wkB332 and wkB2 Δ*tssE1*. (E) Simultaneous colonization of wkB332 and wkB2 Δ*tf*Δ*tssE1*. Fluorescence of GFP (cyan) and E2C (red) are shown in separate and merged channels. Scale bars: 0.5 mm.

### Tailocin and T6SS knockout mutants have no growth rate deficiency

The in vivo effects of mutations affecting tailocin or T6SS potentially reflect changes in the intrinsic growth rate of the mutant strains. To address this, we compared the growth of wkB2 WT and mutants in vitro. wkB2 WT had an exponential doubling time of 1.56 ± 0.02 h (mean ± s.d.). Δ*tf* (1.16 ± 0.03 h) and Δ*tf*Δ*tssE1* (1.20 ± 0.09 h) grew significantly faster than WT, while Δ*tssE1* (1.54 ± 0.06 h) was not significantly different from WT (Fig. 6A). WT reached stationary phase after about 20 h with an optical density at 600 nm (OD_600_) of 0.138 ± 0.006. Δ*tf* (0.354 ± 0.006), Δ*tssE1* (0.308 ± 0.016), and Δ*tf*Δ*tssE1* (0.222 ± 0.013) are significantly higher in maximum OD_600_ at stationary phase after about 40 h (Fig. 6B). Thus, the mutations either confer no difference or confer an advantage in intrinsic growth rate, implying that effects on growth rate cannot explain their impaired competitiveness within the host gut.

**Figure 6.**
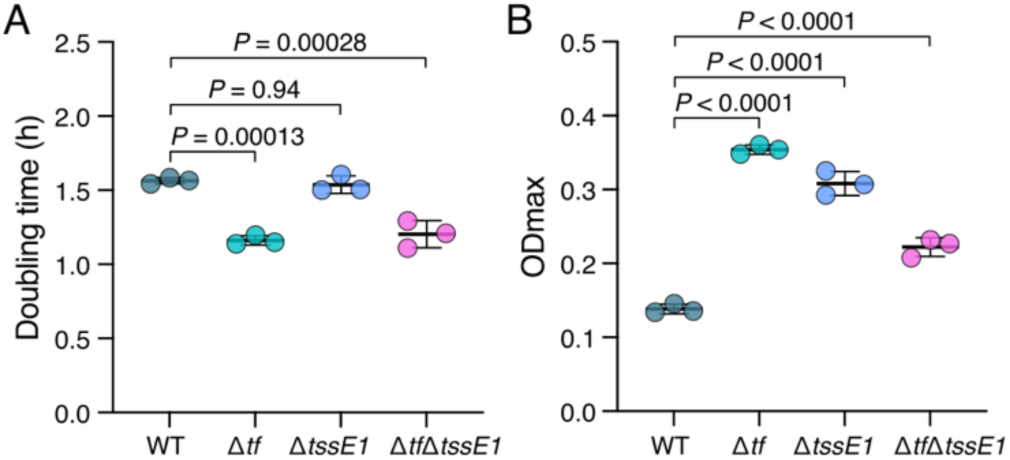
Growth behavior of wkB2 WT and tailocin or T6SS gene deletion mutants. (A) Doubling time of the strains at exponential phase. A one-way ANOVA test (*F*(3, 8) = 40.891; *P* < 0.0001; *n* = 12 biological replicates) indicates significant differences among groups. (B) Maximum optical density (OD) of the strains at stationary phase. A one-way ANOVA test (*F*(3, 8) = 216.4; *P* < 0.0001; *n* = 12 biological replicates) indicates significant differences among groups. Tukey’s Honestly Significant Difference (HSD) test was used for pairwise comparisons of means.

## Discussion

The bee gut microbiome provides a simple system to study strain–strain antagonism in situ. Here, we identified another bacteriocin, tailocin, that contributes to intraspecies competition and investigated its role and interplay with T6SS in gnotobiotic bee guts. Our results show that intact tailocin loci are ubiquitous in *Snodgrassella* strains across different bee hosts (Fig. 1). They are actively expressed in vitro and in vivo, even in those strains such as wkB332 that showed no inhibition against other tested strains. While tailocins of closely related strains can have the same inhibition spectrum, as observed for wkB2 and MS1-3, the overall pattern of susceptibility of strains to a tailocin variant is not determined by phylogenetic distance or host groups (Fig. 2D). Predator strains such as wkB2 can be susceptible to tailocins from other strains, e.g., wkB273. This varied susceptibility within natural communities is likely linked to the low sequence similarity of the tail fiber among tailocin variants. As the tail fiber is responsible for binding receptors on the prey cell surface, its rapid evolution reflects dynamic changes in specificity.

Both weapons are costly, as deletion mutants grow faster and accumulate more cell mass than WT (Fig. 6). The existence of strains maintaining both weapons, such as wkB2, points to intense competition within the bee gut such that the competitive advantage conferred by these weapons outweighs their associated cost. We found that the predator strain wkB2, when possessing either or both tailocin and T6SS, outcompetes the prey strain wkB332 (Fig. 4 and Table 1). In most cases, wkB2 loses its advantage when both weapons are absent. Thus, tailocin, like T6SS (39), directly mediates intraspecies competition in the bee gut. When wkB2 strains colonize first, regardless of the presence or absence of the weapons, they always outcompete wkB332, suggesting another basis for a priority effect independent from tailocin and T6SS. The outcome of competition is heavily influenced by inoculation timing, supporting prior observations of priority effects in bee gut communities (17, 18, 25).

The possession of both weapons does not show an evident advantage over each weapon singly, suggesting that the two weapons might play redundant roles when the prey is susceptible to both. However, our in vitro assays show that strains resistant to tailocin and/or T6SS are present in natural communities (Fig 2D and (62)), so possessing both weapons provides a double safeguard in competition, although it probably also imposes a double cost. The dynamics of weaponry and resistance appear to be highly dynamic, as co-occurring susceptible strains can acquire resistance to toxins, as shown for colibactin (25) and T6SS (62), and the weaponry itself can be horizontally transferred (38). In our system, the rapid evolution of the tail fiber protein suggests arms-race coevolution with target receptor sites on the cell surface.

Though both are derived from phage, T6SS and tailocin show fundamental differences that are expected to affect their utility in different competitive scenarios. T6SS requires cell–cell contact for injection of effectors into neighboring cells, whereas tailocin is released as a diffusible toxin via sacrifice of the lysing host cell (27, 28). Modeling and in vitro assays show that contact-dependent short-range weapons, such as T6SS, are useful for invasion when the predator strain is outnumbered whereas diffusible long-range weapons such as tailocin, are effective only when the predator is abundant (63). These intriguing findings prompted us to compare their effects in vivo. Given that the early colonizer establishes first, our two sequential orders of inoculation of the predator and prey strains represent scenarios of invasion and defense, respectively. Despite the predicted differences, we found that both tailocin and T6SS enable the dominance of the predator strain as an invader, when the prey strain establishes first. As for simultaneous inoculation, the competition outcomes are not significantly different for the two weapon types, indicating similar effects on invasion and simultaneous competition in the gut. The finding that a diffusible and a contact-dependent weapon confer similar advantages in invasion and defense likely reflects the complexity of spatial structure of the gut. Our fluorescence microscopy results provide a spatial context for niche competition in the gut. Both weapons inhibit the establishment of the target strain in the ileum, particularly its aggregation in the anterior ileum (Fig. 5). Since *S. alvi* depends on host-derived nitrogenous wastes and organic acids secreted at the anterior ileum (64), the occupation of this region likely confers an advantage in nutrient acquisition and may explain the preference of *S. alvi* and a *Serratia marcescens* pathogen for the anterior ileum in monoinoculations (25).

When inoculated first, wkB2, whether or not it possesses either weapon, always successfully defends against invasion by wkB332, implying an advantage independent of the two weapons. Although other possible weaponry cannot be ruled out, a potential explanation is a better ability to compete for nutrients. Priority effects through nutrient pre-emption (exploitative competition) are known in other systems (16, 65, 66). Bakkeren *et al.* (67) found that private nutrients, that is, nutrients used only by the cells investing in their acquisition, are required to overcome nutrient blocking for successful invasion. Together, the combination of an advantage in exploitative competition and an interference weapon enables complete displacement by the invader rather than coexistence (67). In our system, wkB332 is often completely outcompeted or displaced by wkB2, but not vice versa, suggesting that wkB2 has advantages through both nutrient and interference competitions.

Tailocins are relatively well studied in *Pseudomonas* species, where in vitro evidence demonstrates their morphology, structure, and mechanisms for killing activity and sensitivity (27, 43, 44, 68, 69). A recent study by Backman *et al.* (70) showed that pathogenic *Pseudomonas* populations colonizing wild populations of *Arabidopsis thaliana* use tailocins to kill closely related competitors, shedding new light on the role and activity dynamics of tailocins in natural communities. They found that genetically diverse strains carry distinct tailocin variants that target variable outer membrane polysaccharides of co-occurring competitors. The same tailocin and outer membrane receptor variants were discovered in historic herbarium samples, suggesting this tailocin-based competitive model has been maintained across time and space. In a different system, Thappeta *et al.* (71) found that tailocins determine competitive outcomes of pathogenic *Xenorhabdus* species in insect hosts. Tailocins produced by different regionally sympatric *Xenorhabdus* strains showed highly varied intraspecies killing profiles, leading to the hypothesis that frequent coinfection of insect hosts by multiple *Xenorhabdus* strains promotes competition and selection (49). To date, aside from these studies, tailocins have been little studied in natural communities. Our study brings new insights into its role in gut environments.

The production of tailocin particles requires lysis of the producing cells and, like phages, usually requires induction (27, 44). Surprisingly, strain wkB2 produces tailocin constantly, even without mutagen induction. The T6SS-null mutant grows better than WT in vitro and outgrows co-inoculated WT in vivo, which is expected as the mutant is relieved of the cost associated with weapon production. However, the tailocin-null mutant almost always colonizes at lower levels than the WT, even though it grows faster and accumulates more cell mass than the WT in vitro. The importance of tailocin in colonizing the bee gut is also evident in the results of a transposon insertion (Tn-Seq) experiment conducted to identify protein-coding genes of *S. alvi* that affect fitness and facilitate host colonization (72). The Tn-Seq results show that the lysin and its neighboring ATPase (locus tags: SALWKB2_0961 and SALWKB2_0962) are essential in culture; whereas the tail fiber chaperone (SALWKB2_0957) is not essential in vitro but is beneficial for gut colonization. Tailocins have been reported to interact with hosts in some other systems, for example, tailocin induces metamorphosis in a marine tubeworm (73). Besides functioning as a weapon for competition, tailocin appears to contribute to colonization ability in the bee guts.

Taken together, we demonstrated the competitive role of tailocin in the bee gut symbiont *Snodgrassella* and investigated its relationship with T6SS in the context of intrastrain interactions. These findings shed new light on intraspecies competition in natural communities. Tailocins have been detected in a variety of bacterial communities, including soft rot in plants, cystic fibrosis infections in humans, and symbiotic *Xenorhabdus* infections in insects, and they have been proposed for use in control of microbiomes, including pathogen exclusion (41, 49, 50, 70, 74, 75). Although many tailocin systems seem to target only closely related bacterial strains, as we found for *Snodgrassella*, several studies have revealed toxicity of tailocins against distantly related bacteria that occupy the same ecological niche (e.g., (76)), implying that tailocins can play a variety of roles in communities. Further studies on tailocin production and specificity will shed light on their roles and evolution in nature and their potential uses in medicine or agriculture.

## Materials and Methods

### Bacterial strains and cultures

Bacterial strains used in this study are listed in SI Appendix, Table S1. *Snodgrassella* strains and other bee gut isolates were routinely cultured on Columbia agar (Becton Dickinson, Franklin Lakes, NJ, USA) supplemented with 5% sheep blood (HemoStat Laboratories, Dixon, CA, USA), except that *Lactobacillus* and *Bombilactobacillus* strains were cultured on Man, Rogosa, and Sharpe (MRS; HiMedia Laboratories, Maharashtra, India) agar. Agar plates were grown in a 35 °C incubator with 5% CO_2_ (Panasonic, Tokyo, Japan), except that *F. perrara* was grown at 35 °C in an anaerobic chamber (Coy, Grass Lake, MI, USA).

### Tailocin purification

Proteinaceous samples that potentially contain tailocins were purified from liquid culture supernatants using an adapted protocol that combines ammonium sulfate precipitation and ultracentrifugation (74, 77). *Snodgrassella* strains were cultured on blood Columbia agar plates for 2-3 d. Cells were inoculated into 10 mL Columbia broth (Alpha Biosciences, Baltimore, MD, USA) in vented tissue culture flasks (Thermo Fisher Scientific, Waltham, MA, USA). After overnight growth in a 35 °C CO_2_ incubator, 1-2 mL cultures were transferred into 2 x 60 mL Columbia broth in vented flasks and grown overnight. Then, induction was carried out by putting 30 mL culture in a 150 mm petri dish and expose under UV for 30 seconds in a biological safety cabinet (1300 Series A2; Thermo Fisher Scientific). The treated cultures were incubated overnight and used for tailocin purification. Culture supernatants were collected by centrifugation (Avanti J-26 XP; Beckman Coulter, Brea, CA, USA) at 8,000 *g* x 30 min, 4 °C, followed by 0.2 µm filtration. The supernatants were put on ice with stirring, and each 100 mL supernatant was slowly added with 65 mL oversaturated ammonium sulfate at the speed of 1 mL per minute. After keeping static at 4 °C overnight, the titrated suspensions were centrifuged at 22,000 *g* x 1 h, 4 °C. Supernatants were removed, and pellets were carefully washed off and resuspended in 1 mL TN50 buffer (10 mM Tris, 50 mM NaCl, pH 7.5). The freshly purified culture supernatant samples were used within two weeks for experiments.

### In vitro inhibition test

We used soft-agar overlay for the inhibition tests (78). Prey strains were grown in Columbia broth (Alpha Biosciences) or MRS broth using vented flasks in a 35 °C CO_2_ incubator for 2 d. A base layer of Columbia/MRS broth with 1.5% agar was poured into petri dishes. Then, a second layer with 0.75% soft agar and prey strains at final OD_600_ of 0.01 were poured. The supernatant samples purified from different strains were 2x or 10x diluted and 5 µL was spotted onto the agar plates with prey strains. Inhibition was checked after 2 d.

### Cryo-EM

*S. alvi* wkB2 tailocin samples for cryo-EM imaging were prepared as aforementioned and incubated with a prey *S. alvi* strain wkB332. wkB332 was grown in Columbia broth (Alpha Biosciences) using a vented flask in a 35 °C CO_2_ incubator for 1 d. Cells were collected at 6,000 *g* x 5 min, 4 °C, and washed with PBS (pH 7.4). wkB2 tailocin sample in TN50 buffer was added to the cell pellet. Cells were resuspended by pipetting gently and put in a culture tube with cap loose and incubated in a 35 °C CO_2_ incubator for 5 h.

Cryo-EM grids were prepared using C-flat™ ultra-flat holey carbon-coated copper grids (2/1 mesh; Electron Microscopy Sciences, Morgantown, PA, USA) and vitrified with a Vitrobot Mark IV (Thermo Fisher Scientific), as previously described (60). Briefly, 3.5 μL of sample was applied to glow-discharged grids maintained at 24 °C and 100% relative humidity. Excess liquid was removed by blotting for 5 s with a blot force of 3, followed by rapid plunge-freezing in liquid ethane. Imaging was performed on a 200 keV Glacios cryogenic transmission electron microscope (Thermo Fisher Scientific) equipped with a Falcon 4 direct electron detector, with automated image acquisition using EPU software (Thermo Fisher Scientific).

### Gene deletion

*S. alvi* wkB2 genes were knocked out by direct transfer of recombineering cassettes through electroporation, as previously described (79). The primers are listed in SI Appendix, Table S2. Q5 High-Fidelity DNA Polymerase (NEB, Ipswich, MA, USA) were used for PCR reactions. Annealing temperatures were as recommended by NEB Tm calculator (https://tmcalculator.neb.com/). PCR products were collected with AxyPrep Mag PCR Clean-up kit (Axygen Biosciences, Union City, CA, USA). The tail fiber deletion cassette (SI Appendix, Table S3) was synthesized at Twist Bioscience (https://www.twistbioscience.com/, South San Francisco, CA, USA) and amplified with the primer pair B2_tfd-f and B2_tfd-r to generate sufficient copies. About 5 mg of the collected PCR product was transferred to late exponential phase cells via electroporation (2 mm cuvette, 3 kV for about 5 ms). Ampicillin-resistant transformants were verified using the primer pair B2_tfd-S-f and B2_tfd-S-r. The *tssE1* gene deletion cassette was amplified using the primer pair B2_tssE-1-f and B2_tssE-1-r and genomic DNA of Δ*tssE1* from a previous study (39) as template. Spectinomycin-resistant transformants were verified using the primer pair

B2_tssE-1-S-f and B2_tssE-1-S-r. We knocked out *tssE1* again to ensure that all mutants have the same genetic background as the WT. This *tssE1* deletion cassette was transferred to Δ*tf* to generate the double knockout mutant. All mutants and the WT were sent for Illumina whole genome sequencing in SeqCenter (https://www.seqcenter.com/, 200 Mbp). The short reads were mapped to the reference genome (GenBank: CP091515) using *breseq* (80) and other mutations are reported in SI Appendix, Table S4.

### Growth measurement

*S. alvi* wkB2 WT and mutants were inoculated into 10 mL Columbia broth (Becton Dickinson) in vented flasks, with three independent cultures as biological replicates for each strain. After 1 d, OD_600_ were measured for late exponential phase cultures by a BioSpectrometer (Eppendorf, Hamburg, Germany) and diluted with Columbia broth to OD_600_ of 0.1. 180 µL Columbia broth and 20 µL OD_600_ of 0.1 culture were added to a 96-well plate (final OD_600_ ∼0.01), with 4 technical replicates for each culture. Absorbance was measured at 600 nm using a Tecan Spark 10M plate reader (Tecan Group, Männedorf, Switzerland) every 2 h for 2 d, with an incubation condition of 35 °C and 5% CO_2_.

### In vivo colonization and competition assay

Microbiota-deprived honeybees were prepared and inoculated as previously reported (25). *S. alvi* strains were cultured in Columbia broth (Becton Dickinson) using vented flasks for 1-2 d. Late exponential phase cells were collected at 6,000 *g* x 5 min, 4 °C, resuspended in PBS to OD_600_ of 1, and mixed with filtrated 1:1 (w/v) sucrose water (v/v, 80:20) immediately before feeding. Newly emerged bees were pooled and randomly assigned to experimental groups, and each bee received 5 µL of culture for each strain via oral feeding. The inoculated bees were individually put in separate 75 mm autoclavable petri dishes (Eisco Labs, Honeoye Falls, NY, USA) supplied with sucrose water (350 µL in 0.5 mL microcentrifuge tubes, with cap open) and irradiated pollen and kept in a 35 °C incubator with ∼80% relative humidity.

The CFU measurement of bee gut samples was adapted from a method used for the *Drosophila* microbiome (81). 5 d after initial inoculation, bees were dissected and whole guts were placed in 96-well PCR plates (Cat. No. 60-0056; Biologix Group, Jinan, Shandong, China) with 0.5 mm zirconia/silica beads (autoclaved; BioSpec Products, Bartlesville, OK, USA) and 100 µL PBS. The PCR plates were sealed with aluminum film (Thermo Fisher Scientific), fixed on a Mini-Beadbeater-96 (BioSpec Products), and homogenized for 45 s. The gut homogenates were serially diluted with PBS and 10 µL were spotted onto blood Columbia agar plates, with 2 technical replicates. Agar plates were cultured in a 35 °C CO_2_ incubator for 3 d when CFUs were counted. We used different antibiotics for selective growth of different strains with different antibiotic markers, as reported before (39). wkB2 Δ*tf*, Δ*tssE1*, Δ*tf*Δ*tssE1*, and wkB332R were selectively grown using ampicillin (50 µg/mL), spectinomycin (30 µg/mL), ampicillin + spectinomycin, and rifampicin (50 µg/mL), respectively. Total abundance was measured using tetracycline (12.5 µg/mL), as wkB2 and wkB332 have natural tetracycline resistance. When wkB2 WT was co-inoculated with another strain, its abundance was calculated by subtracting the other strain from the total abundance (values ≤ 0 were excluded).

### Fluorescence microscopy

Kanamycin-resistant fluorescence plasmids pSL1-GFP (constitutive expression of GFP) and pSL1-E2C (constitutive expression of E2C) (55) were transferred into *S. alvi* wkB332 and wkB2 strains via conjugation as previously reported (82). Strains were grown in Columbia broth (Becton Dickinson) with 25 µg/mL kanamycin to late exponential phase and fed to newly emerged bees as above described. The bees were supplied with sucrose water + 25 µg/mL kanamycin and no pollen (which generates strong autofluorescence). After 5 d, the fluorescence in ilea was inspected via a Nikon Eclipse TE2000-U Inverted Fluorescence Microscope (Nikon Instruments, Melville, NY, USA) and the NIS-Elements AR software (version 5.30.05).

### SDS-PAGE and proteomics

Purified supernatant samples were mixed with a resuspension buffer at final concentrations of 2% SDS, 0.1 M Tris-HCl, and 0.1 M DTT. After treatment of 100 °C for 10 min, an equal volume of 8M urea (pH 8) was added and mixed gently. Then, supernatants were collected after centrifugation at 14000 *g* x 20 min, 4 °C. Protein concentrations were measured by NanoDrop (Thermo Fisher Scientific). 20 µL samples were loaded with 6 x Laemmli buffer into a 12% acrylamide MINI-PROTEAN^®^ TGX™ precast protein gel (Bio-Rad, Hercules, CA, USA) and run with a Tris-glycine-SDS buffer (pH 8.3) at 120 V for about 1 h. A 10-250 kDa prestained protein marker (Cat. No. F4005, APExBIO, Houston, TX, USA), was included. The gel was stained with heated Coomassie Brilliant Blue G-250 (3 g L^−1^ in 45% H_2_O, 45% methanol, 10% acetic acid) for 10 min, then destained overnight in a destain solution (70% H_2_O, 20% methanol, 10% acetic acid).

For proteomics, samples were processed in the same way, except that the loaded samples were run on a 12% acrylamide gel at 100V for ∼10 min until samples migrated out of wells. After staining and destaining, the stained areas were cut into ∼1 mm cubes, in-gel digested with trypsin and desalted with C18 ziptip (Millipore Sigma, Burlington, MA, USA), followed by liquid chromatography with tandem mass spectrometry (LC-MS/MS) analysis on a Vanquish NEO UPLC system coupled with an Orbitrap Ascend Tribrid Mass Spectrometer (both from Thermo Fisher Scientific). Peptides were separated using a 60-min gradient. The raw spectra were analyzed on the Proteome Discover software (version 2.5, Thermo Fisher Scientific) and Scaffold 5 (version 5.3.1, Proteome Software) with default settings (protein threshold, 99.0%; min # peptides, 2; peptide threshold, 95%) and annotated to reference proteomes of respective strains downloaded from UniProt and a list of common contaminants.

For bee-gut-purified samples, wkB2 WT and Δ*tf* were inoculated as aforementioned. The gnotobiotic bees were dissected after 10 d. Ten hindguts were pooled, homogenized with a pestle, added PBS to 1 mL, and vortexed thoroughly. Supernatants of the gut homogenates were collected after centrifugation at 8000 *g* x 5 min, 4 °C, followed by 0.2 µm filtration. 650 µL oversaturated ammonium sulfate was slowly added, vortexed to mix, and left on ice in between. After leaving static at 4 °C overnight, the suspensions were centrifuged at 14,000 *g* x 30 min, 4 °C. Supernatants were removed. Pellets were resuspended in 50 µL TN50 buffer and processed in the same way as above for proteomic sequencing.

### Statistical analysis

The sample sizes were maximized within the constraints of sample handling time. Statistical analyses and data visualization were performed in R (version 4.4.2; http://www.R-project.org/) and R studio (version 2024.09.1; http://www.rstudio.com/). Normal distribution and homogeneity of variances were checked for parametric tests by Shapiro–Wilk test in base R and Levene’s test in the R package car (version 3.1.3) (83), respectively. ANOVA test was conducted using the package car, followed by post-hoc pairwise comparisons of estimated marginal means with a Benjamini-Hochberg adjustment using the package emmeans (84) or of means using Tukey’s Honestly Significant Difference (Tukey’s HSD) in base R. Two-tailed *t*-test was performed in base R. For non-parametric tests, Kruskal–Wallis tests were assessed in base R, followed by two-tailed Dunn’s tests with Benjamini-Hochberg adjustments for post-hoc pairwise comparisons using the package FSA (version 0.10.0) (85). Pearson’s Chi-squared test was assessed in base R, followed by two-tailed Fisher’s exact tests for post-hoc pairwise comparisons using the package rcompanion (version 2.5.2) (86). Package ggplot2 (version 4.0.2) (87) was used for data visualization.

### Bioinformatics

All available *Snodgrassella* genomes were downloaded from the National Center for Biotechnology Information (NCBI). Metagenome-assembled genomes (MAGs) were excluded. Genomes of isolates were quality checked using CheckM (version 1.2.2; lineage_wf mode) (88). Genomes with a completeness of ≤90% or a contamination of ≥5% were excluded. Phage-like sequences were predicted using a standard operating procedure (89) that combines VirSorter2 (version 2.2.4) (90), CheckV (version 1.0.3; database version1.5) (91) and DRAMv (version 1.5.0) (92). The predicted phage-like structures were preliminarily annotated and visualized using Pharokka (version 1.7.3) (93) for manual inspection. The clustering of the phage-like loci was based on pairwise k-mer Mash distances (58) and the elbow method; the analysis was adapted from code made available by T. Backman (59). The PHASTEST web server (https://phastest.ca/) (94) was used for precise annotation of the wkB2 tailocin locus.

*Snodgrassella* genomes were processed using the Genome Taxonomy Database toolkit (GTDB-tk, version 2.4.1) (95) with a database of release 226. A concatenated alignment of 120 bacterial single-copy genes was generated and used to infer a maximum-likelihood tree via IQ-tree 2 (version 2.3.0) (96) The phylogenomic tree and annotations of phage-like clusters were visualized using iTOL (97). Synteny of putative tailocin loci was plotted using clinker (version 0.0.30) (98). The gene map of wkB2 tailocin locus was plotted using Gene Graphics (99).

## Acknowledgments

We thank the Andreas Matouschek lab at UT Austin for use of their ultracentrifuge and William Kern for assistance. We thank Laila Phillips for tips on the CFU assay, and Kim Hammond for maintenance of bee colonies. This study was funded by U.S. National Institutes of Health awards to NAM (R35GM131738 and R35GM158145) and a U.S. Army Research Office grant to Jeffrey Barrick (W911NF-20-1-0195). Cryo-EM images were obtained by an instrument partially supported by the National Institutes of Health (NIH) project number: 1S10OD028700-01. The proteomics analyses were performed at the UT Austin Center for Biomedical Research Support Biological Mass Spectrometry Facility (RRID: SCR_021728).

## Author Contributions

Y.S., N.A.M. conceived the study; Y.S., N.N., N.K., K.R.J., C.L., N.A.M. designed experiments; Y.S., N.N. performed in vitro experiments; Y.S., K.R.J., M.W. performed in vivo experiments; Y.S. analyzed data; Y.S. performed fluorescence microscopy; N.K. performed cryo-EM imaging; N.A.M., C.L. provided resources and supervision; Y.S., N.A.M. wrote an original draft of the ms.; N.N., N.K., CL also contributed to writing and editing. All authors read and gave comments on the ms.

## Competing Interest Statement

The authors declare no competing interest.

## Supporting Information Text

### Identification and clustering of phage-like sequences

275 phage-like loci were predicted in 83 *Snodgrassella* genomes. A manual curation of the annotations and plots of the phage-like structures excluded 44 contigs that only encode some enzymes (e.g., nucleases, methyltransferases, DNA primases and integrases), transcriptional repressors and regulators but no phage structural proteins. The rest 231 loci were partitioned into 5 clusters based on pairwise k-mer Mash distances and the elbow method (1, 2) (Fig S1A). Four clusters (c1, c2, c4, and c5) appear to have complete sets of phage structural proteins (head, baseplate, and tail) and enzymes for replicating and packing were assigned as prophages. One cluster (c3) of 78 contigs that possess tail and baseplate structural proteins but no head proteins and phage enzymes were identified as tailocins. This cluster comprises 3 subclusters (c3.1, c3.2, and c3.3) based on pairwise k-mer Mash distances and the elbow method (Fig S1B).

### Batch-to-batch variation in purification of tailocin-containing supernatant samples and SDS-PAGE

We noticed that supernatant samples purified in different batches differed in protein contents as reflected on the SDS-PAGE gels. In terms of the wkB2 samples, some purification attempts seem to be higher in purity as there are only few major bands on the gel (Fig. 2B); some have numerous extra bands (Fig. S3). They all exhibited normal inhibition against prey strains. The observed differences are likely due to slight differences in growth stage, as the older cultures may release more proteins into the spent medium.

## SI Figures

**Fig. S1.**
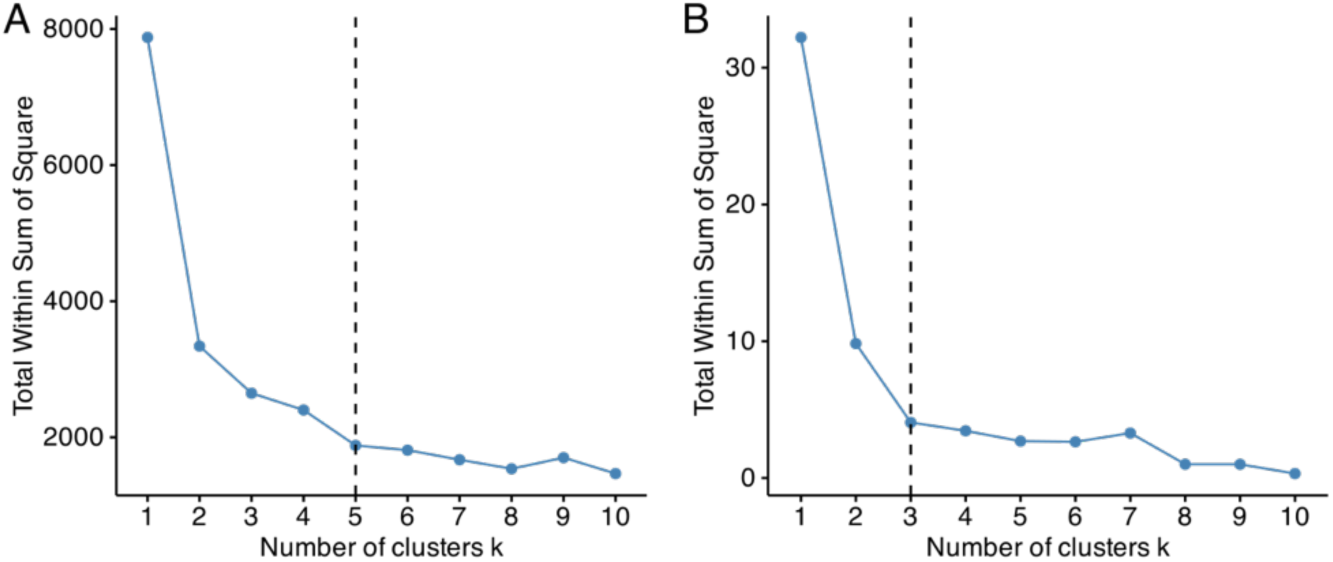
The elbow method indicates that the optimal number of clusters is 5 for all viral-like loci (A) and 3 for predicted tailocin loci (B).

**Fig. S2.**
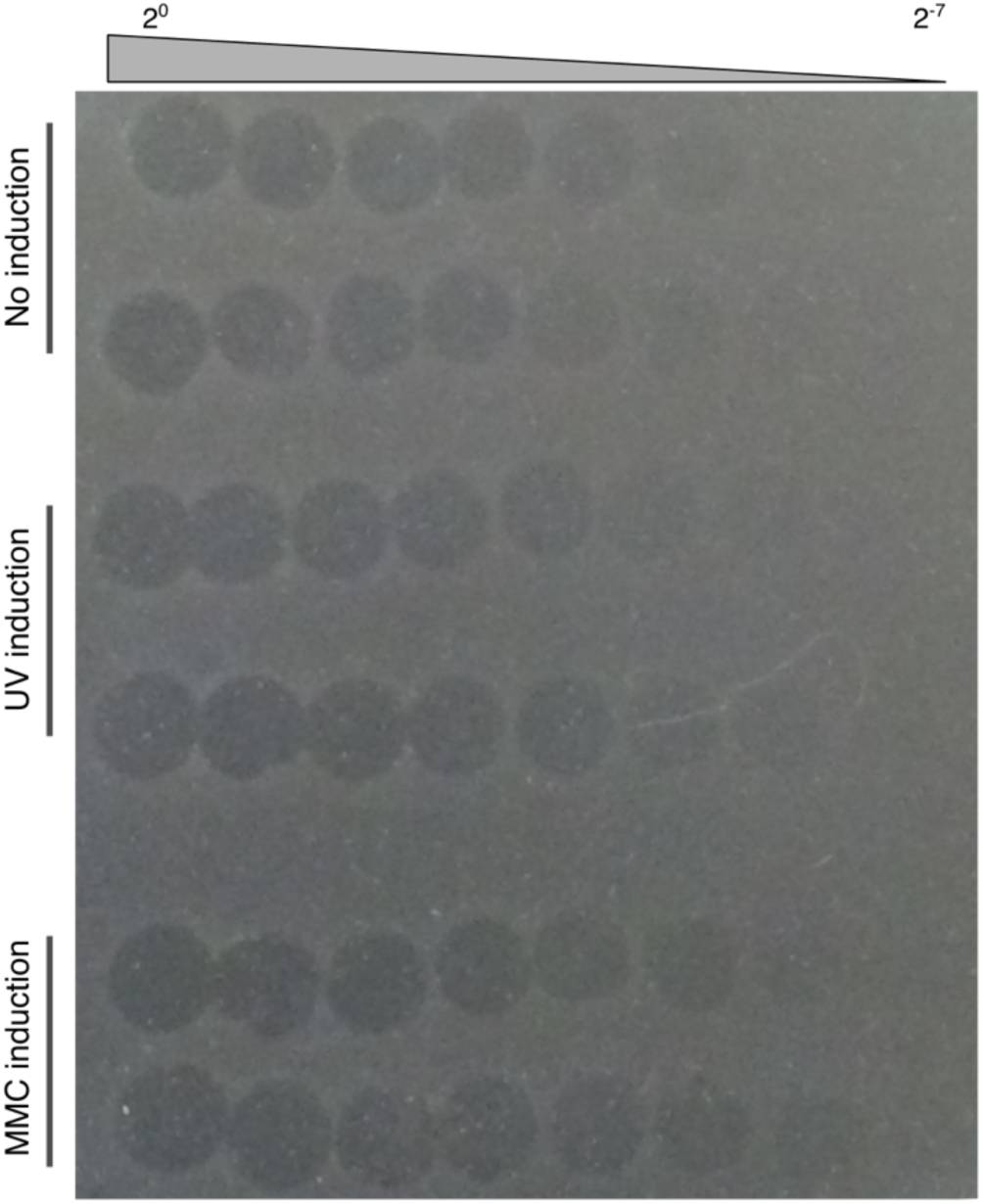
Inhibition of the target strain wkB332 by wkB2 supernatant samples. Samples were purified from wkB2 culture induced by UV, MMC (3 µg/mL), or not induced. Each treatment contains two biological replicates purified from independent cultures.

**Fig. S3.**
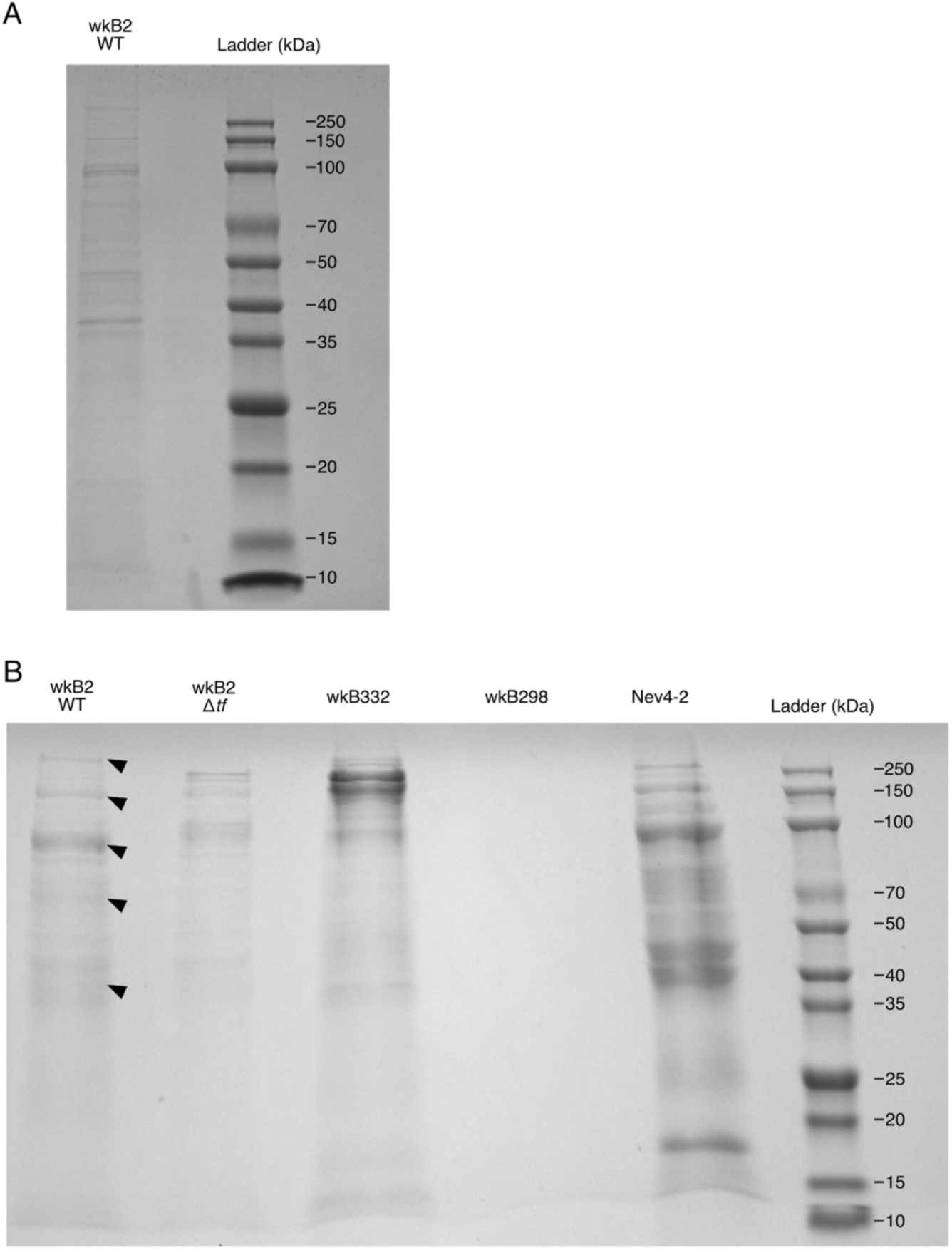
SDS-PAGE analysis of culture-supernatant-purified samples used for proteomic sequencing. (A) A sample purified from strain wkB2 in the first batch (ID: 9871). Arrows show that wkB2 WT has some characteristic bands compared to Δ*tf*. (B) Samples purified from multiple strains in the second batch (ID: 11194). Sample wkB298 was not sequenced as we repeated twice but did not get visible bands for this one.

## SI Tables

**Table S1.** Bacterial strains used in this study.

| Species | Strain | Isolation source | Antibiotic resistance | Reference |
| --- | --- | --- | --- | --- |
| *Snodgrassella alvi* | wkB2 | *Apis mellifera* | Tet | (3) |
| *Snodgrassella alvi* | wkB2 $\Delta tf$ | - | Tet, Amp | This study |
| *Snodgrassella alvi* | wkB2 $\Delta tssE1$ | - | Tet, Spec | (4) and this study |
| *Snodgrassella alvi* | wkB2 $\Delta tf\Delta tssE1$ | - | Tet, Amp, Spec | This study |
| *Snodgrassella alvi* | wkB332 | *Apis mellifera* | Tet | (5) |
| *Snodgrassella alvi* | wkB332R | - | Tet, Rif | (4) |
| *Snodgrassella alvi* | MS1-3 | *Apis mellifera* | - | (5) |
| *Snodgrassella alvi* | PEB0178 | *Apis mellifera* | - | (5) |
| *Snodgrassella* spp. | wkB237 | *Apis andreniformis* | - | (5) |
| *Snodgrassella* spp. | wkB273 | *Apis florea* | - | (5) |
| *Snodgrassella* spp. | wkB298 | *Apis cerana* | - | (5) |
| *Snodgrassella communis* | wkB29 | *Bombus vagans* | - | (6) |
| *Snodgrassella communis* | Occ4-2 | *Bombus occidentalis* | - | (5) |
| *Snodgrassella communis* | Fer1-2 | *Bombus fervidus* | - | (5) |
| *Snodgrassella* spp. | Nev4-2 | *Bombus nevadensis* | - | (5) |
| *Snodgrassella* spp. | App2-2 | *Bombus appositus* | - | (5) |
| *Snodgrassella* spp. | HK3 | *Bombus pensylvanicus* | - | (5) |
| *Gilliamella apicola* | wkB7 | *Apis mellifera* | - | (3) |
| *Gilliamella apis* | M6-3G | *Apis mellifera* | - | (7) |
| *Frischella perrara* | PEB0191 | *Apis mellifera* | - | (8) |
| *Bartonella apis* | PEB0150 | *Apis mellifera* | - | (9) |
| *Bifidobacterium asteroides* | wkB338 | *Apis mellifera* | - | (10) |
| *Lactobacillus helsingborgensis* | wkB8 | *Apis mellifera* | - | (11) |
| *Bombilactobacillus mellifer* | Bin4Na | *Apis mellifera* | - | (12, 13) |

**Table S2.**
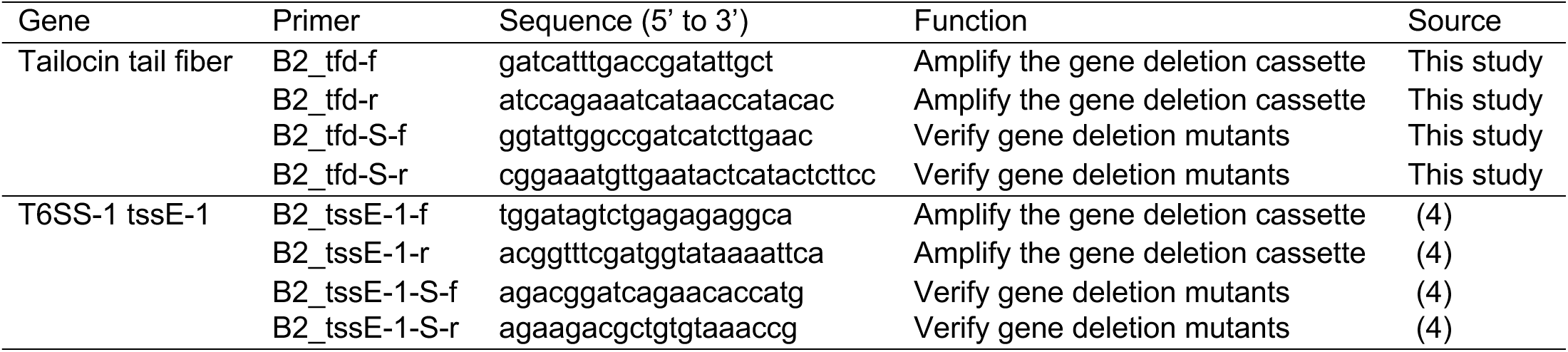
Primers used for *S. alvi* wkB2 gene deletion.

**Table S3.**
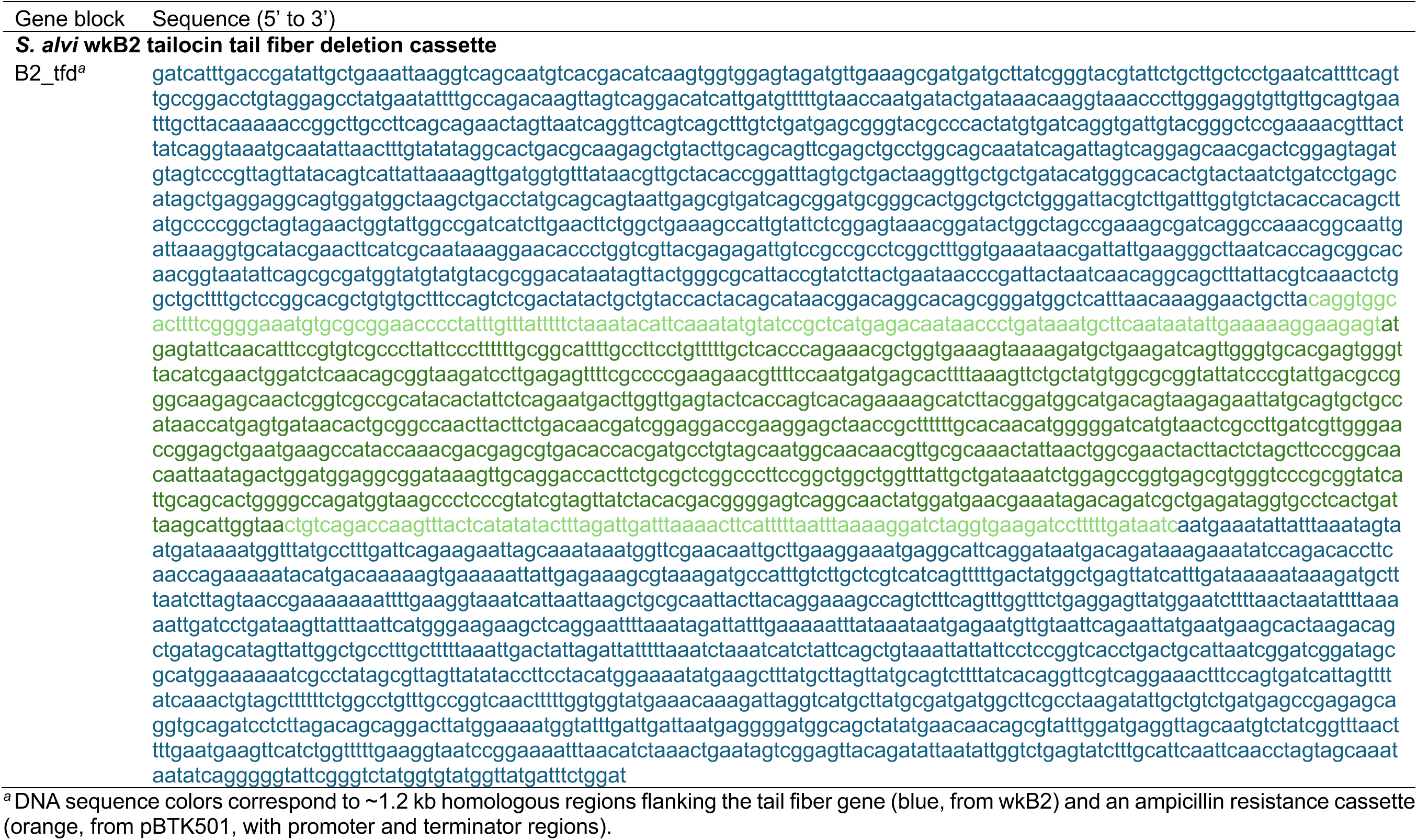
Synthesized DNA fragment of tail fiber gene deletion cassette of *S. alvi* wkB2 tailocin. Gene block Sequence (5’ to 3’)

**Table S4.** Mutations detected in the *S. alvi* wkB2 strains used in this study compared to the reference genome (GenBank: NZ_CP007446.1)

| Strain | Position | Mutation | Gene and product |
| --- | --- | --- | --- |
| wkB2 WT | 1,527,570 | $\Delta 69$ bp<sup>a</sup> | SALWKB2_RS07000 → hypothetical protein |
| wkB2 $\Delta tf$ | 1,527,570 | $\Delta 69$ bp | SALWKB2_RS07000 → hypothetical protein |
|  | 1,058,877 | C→A | SALWKB2_RS04840 → phage tail protein |
|  | 1,058,944 | C→A | SALWKB2_RS04840 → phage tail protein |
| wkB2 $\Delta tssE1$ | 1,527,570 | $\Delta 69$ bp | SALWKB2_RS07000 → hypothetical protein |
| wkB2 $\Delta tf\Delta tssE1$ | 1,527,570 | $\Delta 69$ bp | SALWKB2_RS07000 → hypothetical protein |
|  | 1,058,877 | C→A | SALWKB2_RS04840 → phage tail protein |
|  | 1,058,944 | C→A | SALWKB2_RS04840 → phage tail protein |
|  | 442,282 | T→G | SALWKB2_RS01915 → MFS transporter |

<sup>a</sup> Note that this deletion mutation is inherited from the WT stock and present in all strains.

## Notes

### Competing Interest Statement

The authors have declared no competing interest.

